# *Candida albicans* genome adaptation in azole resistant isolates from autoimmune polyendocrinopathy candidiasis ectodermal dystrophy (APECED) patients

**DOI:** 10.1101/2025.02.17.635002

**Authors:** Faten Alwathiqi, Sara Gago, Lily Novak-Frazer, Riina Richardson, Paul Bowyer

## Abstract

Chronic mucocutaneous candidosis due to *Candida albicans* is frequent in patients with autoimmune polyendocrinopathy candidosis ectodermal dystrophy (APECED). These patients require lifelong antifungal treatment with azoles which might result in the emergence of azole-resistant *C. albicans* isolates. However, the molecular mechanisms that allow *C. albicans* to cause disease and adapt to antifungal treatment in these patients in vastly unknown. Here we comparatively analysed the genome sequences and antifungal susceptibility profile of 14 *C. albicans* isolates from the oral cavities of five APECED patients. 14/13 *C. albicans* isolates showed reduced-susceptibly or resistance to fluconazole. Phylogenetic analyses demonstrated that all APECED isolates from individual patients arose from patient-specific lineages. Single nucleotide variants in genes related to homologous DNA repair mechanisms such as meiosis were associated with APECED (p <0.0001). Altogether our data demonstrates that antifungal adaptation of *C. albicans* in APECED patients might be associated with polymorphisms in proteins responsible for maintaining DNA repair mechanisms.

**Importance:** Chronic mucocutaneous canididosis (CMC) caused by *Candida albicans* is common in patients with primary immunodeficiencies such as autoimmune polyendocrinopathy candidosis ectodermal dystrophy (APECED). These patients receive to long-term antifungal therapy thus promoting the emergence of antifungal resistance. Our knowledge about the mechanisms facilitating CMC in patients with APECED is very limited. Here we demonstrated that most *C. albicans* isolates from patients with CMC are resistant to azoles and that polymorphisms in genes responsible for maintaining DNA repair mechanisms might facilitate infection.

---

Autoimmune polyendocrinopathy candidosis ectodermal dystrophy (APECED) is a long-term autosomal recessive disease caused by loss-of-function mutations of the autoimmune regulator gene (*AIRE*) (Siikala *et al*., 2010) (Moraes-Vasconcelos *et al*., 2008). These mutations result in negative selection if autoreactive T-cells which leads to an autoimmune reaction against endocrine glands and, the development of auto-antibodies against type I interferons (Siikala *et al*., 2010). APECED manifests early in the childhood as a combination of chronic mucocutaneous candidosis (CMC), hypoparathyroidism, and adrenocortical failure (Perheentupa, 2006; Kisand and Peterson, 2015).

CMC in patients with APECED appears first as an oral thrush but as the disease progresses, the whole mouth is involved resulting in difficulties in consumption. *Candida albicans* is the most common *Candida* species causing CMC in these patients (McManus et al., 2011) in which clinical management relies on the use of intermittent therapy with topical and systemic antifungal drugs, mainly polyenes and azoles (McManus *et al*., 2011; Husebye *et al*., 2009). However, long-term antifungal treatment in these patients has previously been linked with the emergence of antifungal resistance (Rautemaa *et al*., 2007b; Rautemaa *et al*., 2007a; Siikala *et al*., 2010). Using gene-targeted approaches, it has previously been reported that point mutations in the target enzyme for azoles (ERG11), TAC1 or overexpression of gene encoding for CDR1/CDR2 and MDR1 efflux-pumps might be linked with persistent *C. albicans* infections in APECED patients (Siikala *et al*., 2010; McManus *et al*., 2011). (McManus *et al*., 2011) but other mechanisms are likely involved.

Here we performed a correlation analyses between azole susceptibility profile and genome composition of 14 *C. albicans* isolates from the oral cavities of five APECED patients from Finland (n = 13) and UK (n = 1). For each patient, at least two *C. albicans* isolates displayed decreased susceptibility to fluconazole or signs of clinical resistance (**Supplementary Table 1 and Figure 1A**). Eleven out fourteen isolates were resistant (> 4 mg/l), two were intermediate and one was susceptible (≤ 2 mg/l) to fluconazole according to EUCAST (ref).

**Figure 1.**
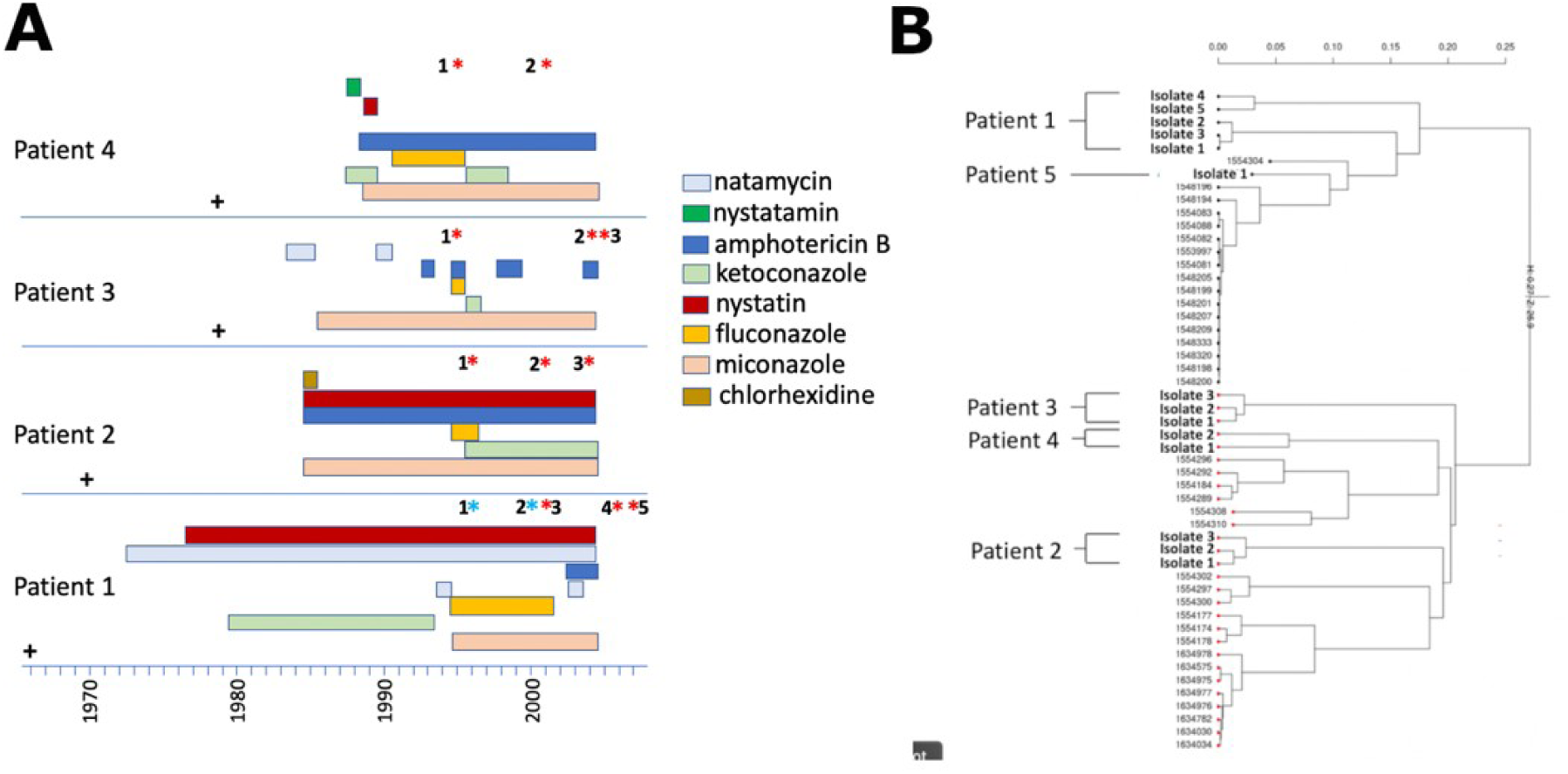
**A.** Treatment timeline and fluconazole-resistance development in *C. albicans* isolated from four APECED patients in Finland. Dates of isolation for all *C. albicans* isolates included in this study are shown as asterisks. Blue asterisks represent fluconazole intermediate isolates and red asterisks represent fluconazole resistant isolates. Treatment courses after 2004 were not available. Details of minimum inhibitory concentrations and dates of recovery are given in Table 1. +; patient date of birth. **B.** Phylogenetic tree of the 14 APECED isolates from five patients and 37 non-APECED isolates from NCBI. Genetic relatedness was calculated using the SNPrelate package in R using identity by descent with 22 genomic regions. The scale bar shows the genetic distance between the isolates.

**Table 1.**
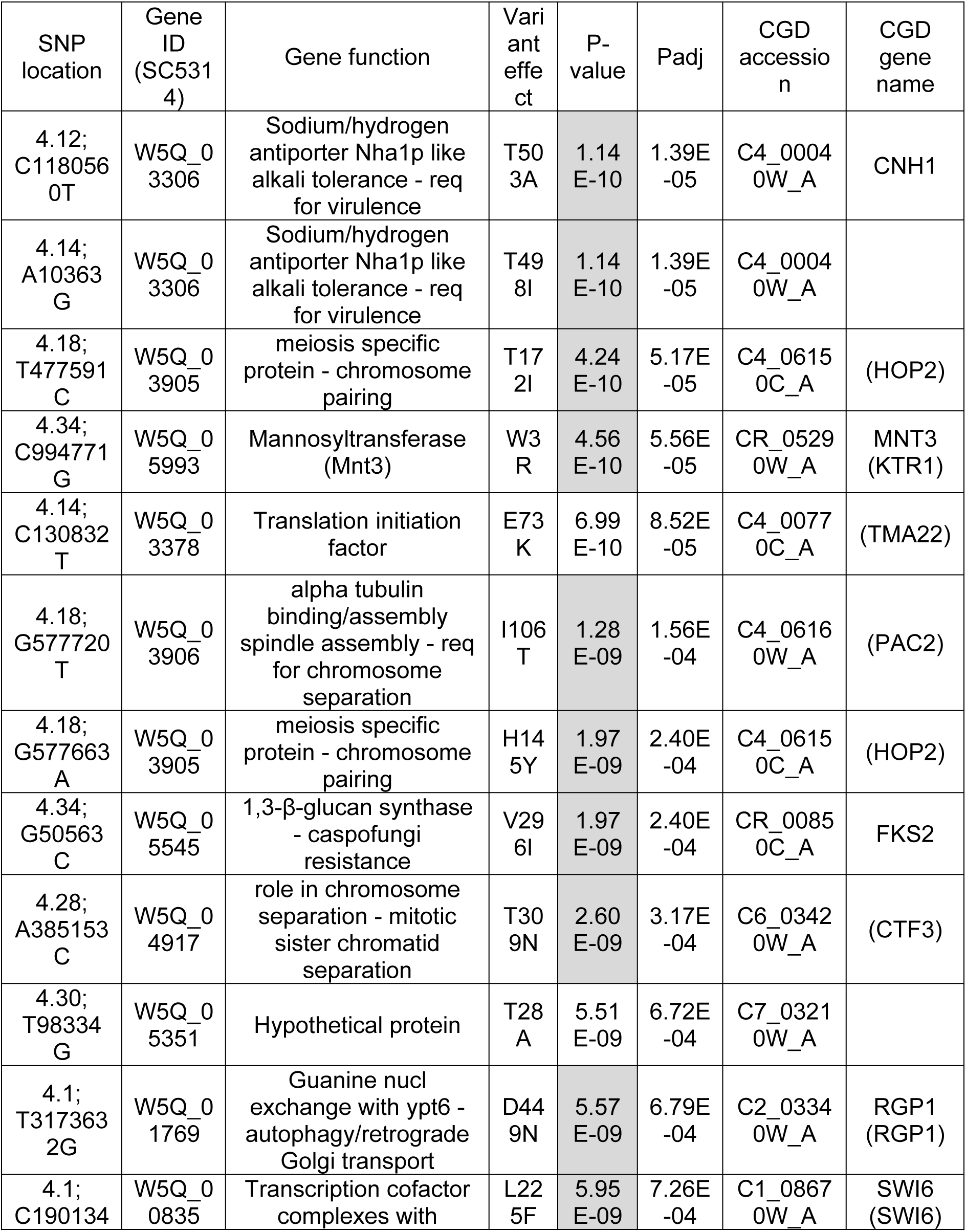
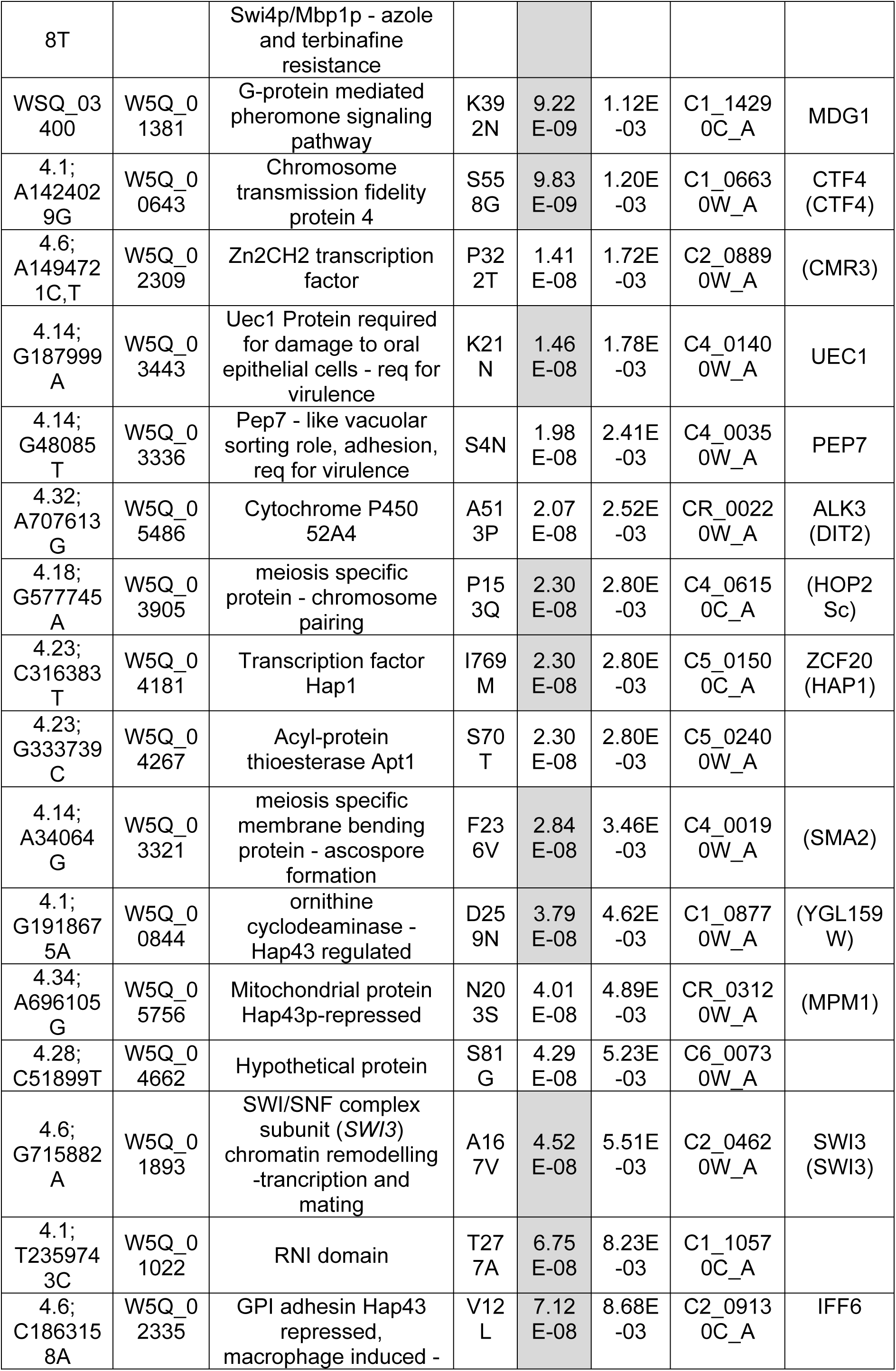
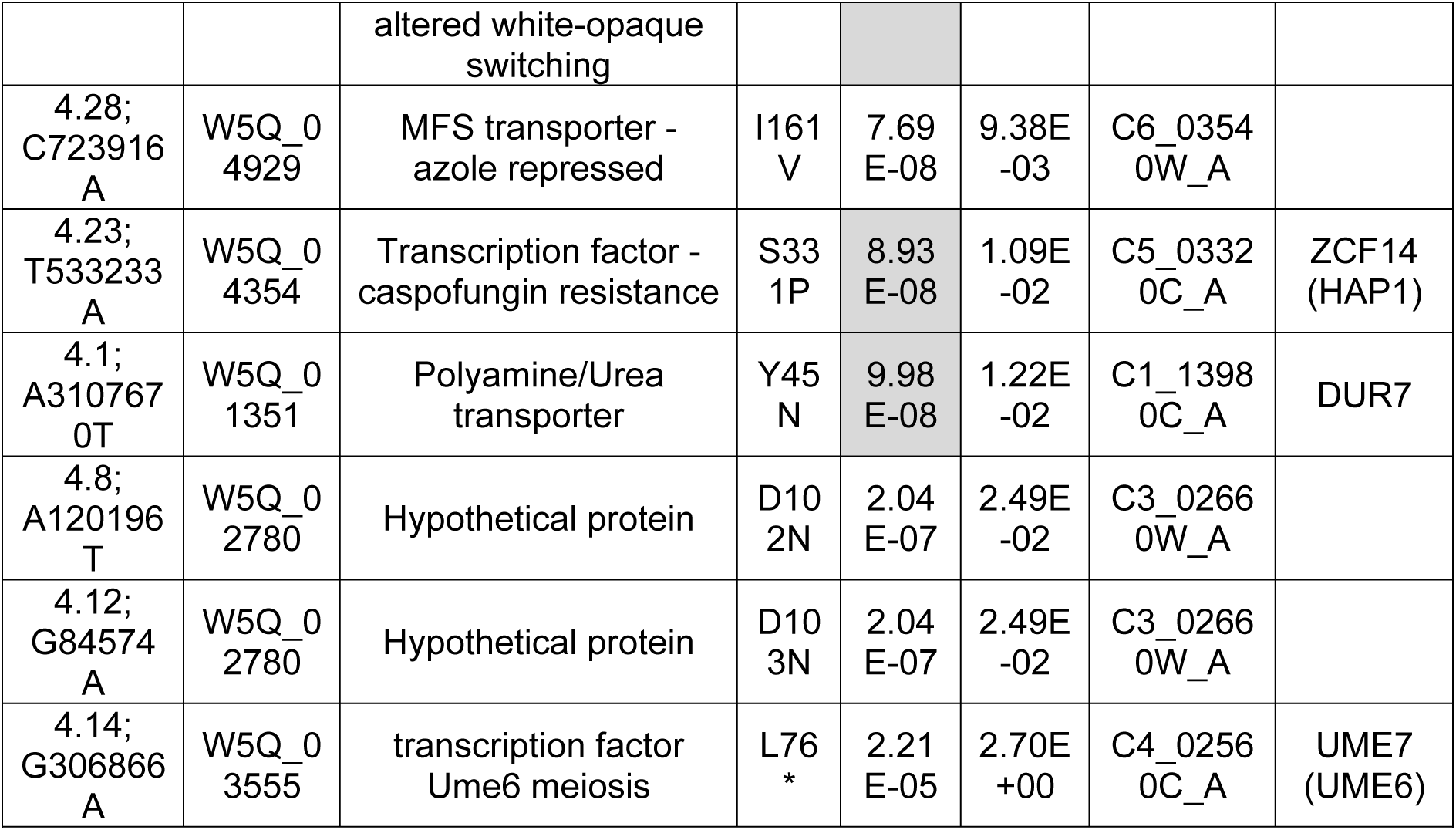
SNPs significantly associated with *C. albicans* isolates from APECED patients. Table shows the location of nucleotide substitution, amino acid change, gene’s ID and function sorted by *Padj*-value. Grey boxes indicate P values supported by lineage-specific analysis in ROADTRIPS. *Candida* gene names are listed where available and *Saccharomyces cerevisiae* orthologue names are given in brackets

We next explored whether the *C. albicans* isolates included in this study were phylogenetically related by using whole genome sequencing. Our data indicates that *C. albicans* isolates from different APECD patients belong to different lineages. However, *C. albicans* isolates within each patient are lineage-specific with the exception of patient 1 where independent but related lineages were shown (**Figure 1B**).

To investigate whether genetic variability in *C. albicans* contributes to APECED, we performed a global analyses of genetic polymorphisms in all 14 APECED isolates included in the study and 54 non-APECED *C. albicans* controls. All genomes were aligned to the *C. albicans* SC5314 reference genome. A total of 155661 raw variants were found but this figure was reduced to 84314 variants when variation arising from reference genome sequence version was accounted for. SNPs were further filtered to remove MAF <0.01, deviation from strand >0, presence in the dataset <0.9. After filtering 35 variants with high moderate or low impact as assessed using SNPEff remained **(Table 1).** Ten of the discovered variants were functionally associated with meiosis (Padj <0.0001) and sporulation and, 8 of them were specifically associated with chromosome pairing, segregation or sister chromatid exchange in homologous DNA repair pathways (Padj <0.0001). Only 4/35 discovered variants had functional roles in virulence (FET < 0.05). This is not surprising given the wide assignation of virulence phenotypes (967/6137 genes) and the fact that virulence is assessed using immunocompromised mouse models which are likely to predict APECED poorly.

Since most *C. albicans* APECED isolates were resistant to fluconazole, we also assessed genetic variants in known genes linked with azole resistance (ERG11, CDR1, CDR2, TAC1, MDR1 and MRR1). We found that 7/10 azole resistant isolates tested carried previously known resistance variants (Siikala *et al*., 2010; McManus *et al*., 2011) in ERG11, and 4 of them also had known resistance-associated variants in TAC1 with no comparable variants found in 54 azole sensitive, non-APECED controls.

Two additional variants were in genes with possible association to azole resistance (SWI6 and C6_03540W_A) but the link appears tenuous (p values??). Even though the APECED patients had no previously received echinocandin treatment, we found three variants in genes with strong links to caspofungin resistance (FKS2, ZCF20 and ZCF14) in these patients.

Copy number variation in genes involved in antifungal resistance can trigger changes in expression of target enzymes and the development of antifungal resistance (ref??). We only found one isolate (Patient 1, isolate 1) carrying a duplication consisting of a 76 gene region (SC5314 supercont4.1: 2,162,420-2,360,571) (**Supplementary Table 2)**. Even though the duplicated region contained several genes involved in cell wall formation and immune system evasion (MNN2 and MNN12) and a range of transcription factors notably WOR1, the deleted region did not contain any genes linked with azole resistance. No enrichment for any functional group of genes was observed in this region.

Here, we presented for the first time a comparative analyses of the genomes of *C. albicans* isolates from patients with long-term APECED. In agreement with previous studies, our results indicate that isolates from APECED patients are from clonal lineages (McManus, Moorhouse). *C. albicans* APECED isolates carried SNPs in genes contributing to mismatch repair and double-strand break repair that have been previously linked to genomic instability and increased frequency of drug-resistance in both *C. albicans* (Legrand *et al*., 2007) and *C. glabrata*, (Healey et al., 2016). Given lack of any evidence for sexual recombination in our isolates or in other studies of this kind (Moorhouse, McManus) we suggest that the significant enrichment of meiosis type homologous pairing and repair genes observed in APECED isolates may reflect a similar evolution to a mutating phenotype. It is important to note that the patients included in this study have complex treatment histories and therefore results can only represent genome adaptation to the complex APECED stress environment which includes a variety of antifungal drugs, the host immune response and interplay with the oral? microbiome.

## Materials and methods

*Candida albicans* isolates and antifungal susceptibility testing: 14 *C. albicans* isolates from APECD patients from Finland and UK were included in the studyxxx. AFST to fluconazole was carried out as described by the European Committee on Antimicrobial Susceptibility Testing (EUCAST) (Arendrup *et al*., 2015). Patient treatment history is summarised in Figure 1.

Genome sequencing and bioinformatics: **DNA extraction was carried out using the Qiagen Pathogen Lysis tubes and the Qiagen QIAamp DNA mini kit (Qiagen, Germany) following the manufacturer’s instructions. 10 µg of DNA was sequenced using the** Illumina **MiSeq platform (Illumina, USA) in the Genomic Technologies Core Facility at the University of Manchester. The maximum read length was 2 x 250 bp paired-end reads. Sequences were converted to FASTQ format using bcl2fastq v2.17.1.14 (Illumina, USA) to remove any residual adapter sequences.** The reads were checked using FastQC v0.11.3 and fastq-screen v0.5.2 (Bioinformatics, 2015). Further filtering for the per base calling quality and read length was applied using Trimmomatic v0.36 using TRAILING and LEADING Phred score of 30 and a SLIDING WINDOW of 2 bp at a Phred score of 30 (Bolger *et al*., 2014). Secondly, the reads were aligned to *C. albicans* SC5314 (GCA_000784635_V4) using Burrows-Wheeler Alignment tool (BWA-mem) alignment tool v0.7.15-r1140 (Li and Durbin, 2009; Ensembl, 2017). SNP calling was performed using the genome analysis toolkit (GATK 3.7.0) using two rounds of base score quality realignment (McKenna *et al*., 2010). The resulted variant call format (VCF) file was then used to generate an identity-by-descent (IBD) phylogenetic tree using the R package SNPRelate. Also, the SnpEff/SNPSift package was used to assess the impact of the observed variants on protein function (Cingolani *et al*., 2012b; Cingolani *et al*., 2012a). Association of variants with APECED was assessed using 54 other non-APECED *C. albicans* reference isolates as controls (NCBI SRA archive). Fishers exact tests were calculated and Bonferroni correction for multiple sampling was applied. We noted that many SNPs were present in alternative SC5314 references and filtered these sites and variants with low mapping quality from the analysis, leaving 84,000 sites for analysis. The resulting SNPs were categorised using SNPSift/SNPEff. SNPs surviving filters and with impact low, moderate or high were further analysed manually to annotate and assess function using the Candida Genome Database (Table 1). Additionally the strong lineage structure observed in the phylogenetic reconstruction suggested that results should be analysed using an algorithm that allowed for lineage specific effects. SNPs were analysed using lineage specific algorithm RIADTRIPSPolymorphisms in known genes associated with azole-resistance that have high or moderate impact including *ERG11*, *CDR1*, *CDR2*, *TAC1*, *MDR1* and *MRR1* were also collected (Supplementary Table 3).

Copy number variation (CNV) determination: MiSEQ reads were quality filtered as described above then mapped to the entire *C. albicans* SC5314 reference genome split into 500 bp overlapping segments or against cDNAs using bowtie 2 (Cingolani *et al*. 2012) then piped directly through SAM format into a bespoke read counting script. (Li *et al*., 2009). Count per gene or feature was then normalised against SC5314 read counts and against counts per genome to derive a measure of copy number for each gene.

Statistical analysis: Fishers exact tests were performed on all variant calling results using SNPSift (version 4.0). Multiple testing corrections were performed on the Fisher exact results using a Bonferroni correction to exclude any false rate of *P*-values.

---

**Supplementary table 1.** Isolates used in this study. Fluconazole-susceptibility (mg/l) results of the 14 APEPCED isolates.

| Patient | Isolate | Strain number | Date of collection | Fluconazole resistance (MIC, mg/l) |
| --- | --- | --- | --- | --- |
| 1 | 1 | 826 | 1996 | I (4) |
|  | 2 | 109 | 2000 | I (3) |
|  | 3 | 355 | 2001 | R (32) |
|  | 4 | 1382 | 2006 | R (8) |
|  | 5 | 1527 | 2007 | R (32) |
| 2 | 1 | 931 | 1996 | R (16) |
|  | 2 | 366 | 2001 | R (64) |
|  | 3 | 916 | 2004 | R (32) |
| 3 | 1 | 719 | 1995 | R (16) |
|  | 2 | 1108 | 2004 | R (64) |
|  | 3 | 1179 | 2005 | R (32) |
| 4 | 1 | 695 | 1995 | R (32) |
|  | 2 | 373 | 2001 | R (32) |
| 5 | 1 | 56388 | 2015 | S (2) |

**Supplementary Table 2.** List of genes in the *C. albicans* duplicated chromosomal region of Patient 1 isolate 1.

| Gene ID<br>(SC5314) | CGD<br>accession | Gene function | CGD gene<br>name |
| --- | --- | --- | --- |
| W5Q_009<br>46 | C1_09800<br>_A | Golgi apparatus membrane protein (*TVP18*) | TVP18<br>(TVP18) |
| W5Q_009<br>47 | C1_09810<br>_A | Ygr210 GTPase |  |
| W5Q_009<br>48 | C1_09830<br>_A | Superkiller protein 3 | SKI3 (SKI3) |
| W5Q_009<br>49 | C1_09840<br>_A | role in proteasome localisation | STS1<br>(STS1) |
| W5Q_009<br>50 | C1_09850<br>_A | Hypothetical protein |  |
| W5Q_009<br>51 | C1_09860<br>_A | Transcription initiation factor TFIIE subunit $\alpha$ | TFA1<br>(TFA1) |
| W5Q_009<br>52 | C1_09870<br>_A | Forkhead transcription factor (*HCM1*) Hap43<br>induced | HCM1<br>(HCM1) |
| W5Q_009<br>53 | C1_09880<br>_A | Bud site selection protein 31 | BUD31<br>(BUD31) |
| W5Q_009<br>54 | C1_09890<br>_A | tRNA Ser (CGA) |  |
| W5Q_009<br>55 | C1_09900<br>_A | Secreted Lipase 3 | LIP3 |
| W5Q_009<br>56 | C1_09910<br>_A | Phosphatase - Hap43p-induced |  |
| W5Q_009<br>57 | C1_09920<br>_A | Vacuole organisation | VPS41 |
| W5Q_009<br>58 | C1_09930<br>_A | Hypothetical protein |  |
| W5Q_009<br>59 | C1_09940<br>_A | Hypothetical protein |  |
| W5Q_009<br>60 | C1_09950<br>_A | Golgi membrane protein (*TVP23*) | (TVP23) |
| W5Q_009<br>61 | C1_09960<br>_A | Cytochrome C chaperone | PET100 |
| W5Q_009<br>62 | C1_09970<br>_A | Transcription factor(*PDC2*) | PDC2 |
| W5Q_009<br>63 | C1_09980<br>_A | Acyl glycerol lipase | YJU2 |
| W5Q_009<br>64 | C1_09990<br>_A | phosphatidylinositol phosphate (PtdInsP)<br>phosphatase, involved in cell wall integrity and<br>morphogenesis | SAC1<br>(SAC1) |
| W5Q_009<br>65 | C1_10000<br>_A | MAPK phosphatase - roles in filamentation<br>virulence | CPP1<br>(MSG1) |
| W5Q_009<br>66 | C1_10010<br>_A | G2/M protein kinase req for virulence,<br>caspofungin sensitive | SWE1 |
| W5Q_009<br>67 | C1_10020<br>_A | Fe response transcription factor Hap43<br>repressed | SFU1<br>(GLN3) |
| W5Q_009<br>68 | C1_10030<br>_A | ATP-dependent RNA helicase *DBP3* Hap43<br>induced | DBP3 |
| W5Q_009<br>69 | C1_10040<br>_A | Endoplasmic Reticulum Oxidoreductin 1<br>(*ERO1*) | ERO1<br>(ERO1) |
| W5Q_009<br>70 | C1_10050<br>_A | Protein not essential for viability |  |
| W5Q_009<br>71 | C1_10060<br>_A | Biofilm-induced protein |  |
| W5Q_009<br>72 | C1_10070<br>_A | α-1,2-mannosyltransferase amB resistance,<br>ergosterol intermediate resistance | MNN2 |
| W5Q_009<br>73 | C1_10080<br>_A | Pre-mRNA-splicing factor (CWC21) | CWC21 |
| W5Q_009<br>74 | C1_10090<br>_A | Hypothetical protein |  |
| W5Q_009<br>75 | C1_10100<br>_A | Nuclear transport factor 2 |  |
| W5Q_009<br>76 | C1_10110<br>_A | Biofilm-induced protein |  |
| W5Q_009<br>77 | C1_10120<br>_A | Transcription initiation factor TFIIE subunit $\beta$ | |
|  | C1_10130<br>_A | snoRNA |  |
| W5Q_009<br>78 | C1_10140<br>_A | F-box protein COS111 |  |
| W5Q_009<br>79 | C1_10150<br>_A | Transcription factor master switch opaque-<br>white switching | WOR1 |
| W5Q_009<br>80 | C1_10160<br>_A | Mitochondrial phosphate carrier protein | MIR1 |
| W5Q_009<br>81 | C1_10170<br>_A | Putative adhesin-like protein |  |
| W5Q_009<br>82 | C1_10180<br>_A | regulator of endocytosis | ECM21 |
| W5Q_009<br>83 | C1_10190<br>_A | RNA recognition motif |  |
| W5Q_009<br>84 | C1_10200<br>_A | Major Facilitator Superfamily | HOL1<br>(HOL1) |
| W5Q_009<br>85 | C1_10210<br>_A | Serine/threonine-protein kinase req for<br>virulence | CLA4<br>(CLA4) |
| W5Q_009<br>86 | C1_10220<br>_A | cAMP-dependent protein kinase type 3 | TPK1 |
| W5Q_009<br>87 | C1_10230<br>_A | Protein not essential for viability |  |
| W5Q_009<br>88 | C1_10240<br>_A | Ribonuclease activity regulator RraA family |  |
| W5Q_009<br>89 | C1_10250<br>_A | Hypothetical protein |  |
| W5Q_009<br>90 | C1_10260<br>_A | DNA-directed RNA polymerase I subunit<br>(RPA34) | RPA34 |
| W5Q_009<br>91 | C1_10270<br>_A | Phosphatidylinositol transfer protein (SFH5) | SFH5 |
| W5Q_009<br>92 | C1_10280<br>_A | Protein FMP52, mitochondrial |  |
| W5Q_009<br>93 | C1_10290<br>_A | Glucoamylase 1 | GCA1 |
| W5Q_009<br>94 | C1_10300<br>_A | α-1,3-mannosyltransferase | MNN12 |
| W5Q_009<br>95 | C1_10310<br>_A | Methyltransferase domain |  |
| W5Q_009<br>96 | C1_10320<br>_A | N-acetylglucosaminylphosphatidylinositol<br>deacetylase |  |
| W5Q_009<br>97 | C1_10330<br>_A | Actin associated - MMS-induced unequal sister<br>chromatid recombination | ARP6 |
| W5Q_009<br>98 | C1_10340<br>_A | MFS transporter |  |
| W5Q_009<br>99 | C1_10350<br>_A | Adhesin with role in cell cycle |  |
| W5Q_010<br>00 | C1_10360<br>_A | Hypothetical protein |  |
| W5Q_010<br>01 | C1_10370<br>_A | Hypothetical protein |  |
| W5Q_010<br>02 | C1_10380<br>_A | serine/threonine kinase - cell wall integrity | CBK1 |
| W5Q_010<br>03 | C1_10390<br>_A | 60S ribosomal protein L44 |  |
| W5Q_010<br>04 | C1_10400<br>_A | Putative GPI-anchored adhesin-like protein<br>(Fgr41p) | FGR41 |
| W5Q_010<br>05 | C1_10410<br>_A | DNA binding protein |  |
| W5Q_010<br>06 | C1_10420<br>_A | ubiquitin protein ligase binding activity |  |
| W5Q_010<br>07 | C1_10430<br>_A | Putative repressible vacuolar alkaline<br>phosphatase | PHO8 |
| W5Q_010<br>08 | C1_10440<br>_A | proposed role in Golgi vesicle trafficking | (GTA1) |
| W5Q_010<br>09 | C1_10450<br>_A | L-threonine aldolase | GLY1 |
| W5Q_010<br>10 | C1_10460<br>_A | Hypothetical protein |  |
| W5Q_010<br>11 | C1_10470<br>_A | Small subunit ribosomal protein S5 |  |
| W5Q_010<br>12 | C1_10480<br>_A | Uncharacterized ORF (Djp1p) | DJP1 |
| W5Q_010<br>13 | C1_10490<br>_A | Leukotriene A-4 hydrolase | LKH1 |
| W5Q_010<br>14 | C1_10500<br>_A | Hypothetical prtein |  |
| W5Q_010<br>15 | C1_10510<br>_A | Hypothetical protein |  |
| W5Q_010<br>16 | C1_10520<br>_A | Leucine-rich repeat (LRR) protein |  |
| W5Q_010<br>17 | C1_10530<br>_A | Ribosome-associated protein | SNL1 |
| W5Q_010<br>18 | C1_10540<br>_A | Putative adapter protein; links synaptojanins |  |
| W5Q_010<br>19 | C1_10550<br>_A | Glucoamylase 1, partial | GCA2 |
| W5Q_010<br>20 | C1_10560<br>_A | Hypothetical protein |  |
| W5Q_010<br>21 | C1_10570<br>_A | Hap43p-repressed protein |  |
| W5Q_010<br>22 | C1_10580<br>_A | Hypothetical protein |  |

**Supplementary Table 3.** Discovered genetic variants in genes previously known to be involved in azole resistance across all 14 *C. albicans* isolates included in the study. Mutations A114V, Y132F, K143I, D153E, E266D, G307S, V437I, G450E, S405F, R467K and G464S, V488I (in bold) have been previously described to be associated with azole-resistance (Akins and Sobel, 2017; Flowers *et al*., 2015). Isolates 1 and 2 had intermediate fluconazole-susceptibility, patient 1 isolates 3-5 and all isolates from patients 2 and 3 were azole-resistant and patient 5 isolate 1 was azole-susceptible. Mutations R673Q, A736V, N972S, N972D, G980E and E461K (in bold) have been previously described to be associated with azole-resistance (Morschhäuser, 2010; Kalkandelen and Doluca, 2015; Coste *et al*., 2007). Mutation N740S, N977D, P276L, E461K and R673Q (underlined) have been previously identified in the same isolates studied by (Siikala *et al*., 2010).

| Patient |  | ERG1<br>1 | CDR1<sup>a</sup> | CDR2 | TAC1<sup>a</sup> | MDR<br>1<sup>b</sup> | MRR1<sup>c</sup> |
| --- | --- | --- | --- | --- | --- | --- | --- |
| 1 | 1 | E116<br>D,<br>K128<br>T |  | A59T, E179K,<br>Y425F, F439Y,<br>A620T, R683K,<br>P1305L,<br>L1337F,<br>G1473A, I1474V | V104F, N199S,<br>H206R, A207V,<br>N396S, L658F,<br><u>N740S</u>, H741Y,<br>N772K, D776N,<br>Q829E, S935*,<br>L941P | I285T | L171P,<br>L248V,<br>V341E,<br>L592F,<br>E1020Q |
|  | 2 | E116<br>D,<br>K128<br>T |  | A59T, E179K,<br>A620T, R683K,<br>P1305L,<br>L1337F,<br>G1473A, I1474V | P28S, V104F,<br>N199S, H206R,<br>A207V, N396S,<br>L658F, <u>N740S</u>,<br>H741Y, N772K,<br>D776N, Q829E,<br>S935*, L941P | I285T | V27I,<br>L171P,<br>L248V,<br>V341E,<br>L592F,<br>E1020Q |
|  | 3 | E116<br>D,<br>K128<br>T |  | A59T, E179K,<br>Y425F, F439Y,<br>A620T, R683K,<br>P1305L,<br>L1337F,<br>G1473A, I1474V | V104F, S108N,<br>N199S, H206R,<br>A207V, N396S,<br>L658F, <u>N740S</u>,<br>H741Y, N772K,<br>D776N, Q829E,<br>S935*, L941P | I285T | V27I,<br>L171P,<br>L248V,<br>V341E,<br>L592F,<br>E1020Q |
|  | 4 | E116<br>D | A833T | A59T, E179K,<br>V498F | V104F, N199S,<br>H206R, A207V,<br><u>P276L</u>, Q829E,<br>S935*, L941P,<br>**N972S**, <u>N977D</u> |  | V27I,<br>L171P,<br>L248V,<br>V341E,<br>L592F,<br>E1020Q |
|  | 5 | E116<br>D | A833T | A59T, E179K,<br>V498F | V104F, N199S,<br>H206R, A207V,<br>Q829E, S935*,<br>L941P, **N972S**,<br>N977D |  | V27I,<br>L171P,<br>V341E,<br>L592F,<br>E1020Q |
| 2 | 1 | E116<br>D,<br>**D153**<br>**E**,<br>**E266**<br>**D** | E1416<br>A | G50A, Y98C,<br>V121I, E179K,<br>Y425F, F439Y,<br>E445Q, G466R,<br>R470C, A620T,<br>R683K, V767I,<br>A768G, T840S,<br>D1102N,<br>R1144K,<br>L1337F,<br>G1473A, I1474V | V104F, S108N,<br>M170I, F189S,<br>A377V, N396S,<br>I558V, L695F,<br>N772K, D776N,<br>Q829E, N896S,<br>L941P | I285T<br>,<br>N362<br>S,<br>S405<br>T,<br>T634I | L171P,<br>V341E,<br>L592F,<br>E1020Q |
|  | 2 | E116<br>D,<br>**D153**<br>**E**,<br>**E266**<br>**D**,<br>**G450** | E1416<br>A | G50A, Y98C,<br>V121I, E179K,<br>Y425F, F439Y,<br>E445Q, G466R,<br>R470C, A620T,<br>R683K, V767I,<br>A768G, T840S, | V104F, S108N,<br>M170I, F189S,<br>A377V, N396S,<br>I558V, L695F,<br>N772K, D776N,<br>Q829E, N896S,<br>L941P | I285T<br>,<br>N362<br>S,<br>S405<br>T,<br>T634I | L171P,<br>V341E,<br>L592F,<br>E1020Q |
|  |  | **E** |  | D1102N,<br>R1144K,<br>L1337F,<br>G1473A, I1474V |  |  |  |
|  | 3 | **A114**<br>**V,**<br>**D153**<br>**E,**<br>**E266**<br>**D,**<br>**G450**<br>**E** | E1416<br>A | G50A, Y98C,<br>V121I, E179K,<br>Y425F, F439Y,<br>E445Q, G466R,<br>R470C, A620T,<br>R683K, V767I,<br>A768G, T840S,<br>D1102N,<br>R1144K,<br>L1337F,<br>G1473A, I1474V | V104F, S108N,<br>M170I, F189S,<br>A377V, N396S,<br>I558V, L695F,<br>N772K, D776N,<br>Q829E, N896S,<br>L941P | I285T<br>,<br>N362<br>S,<br>S405<br>T,<br>T634I | L171P,<br>V341E,<br>L592F,<br>E1020Q |
| 3 | 1 | E116<br>D,<br>**S405**<br>**F** |  | A59T, E179K,<br>A620T, R683K,<br>P1305L,<br>L1337F,<br>G1473A, I1474V | N396S, **E461K**,<br>D776N, N896S,<br>L941P |  | V27I,<br>L171P,<br>L248V,<br>V341E,<br>L592F,<br>E1020Q |
|  | 2 | E116<br>D,<br>**S405**<br>**F** |  | A59T, E179K,<br>A620T, R683K,<br>P1305L,<br>L1337F,<br>G1473A, I1474V | N396S, **E461K**,<br>D776N, N896S,<br>L941P | P255<br>L | V27I,<br>L171P,<br>L248V,<br>V341E,<br>L592F,<br>E1020Q |
|  | 3 | E116<br>D,<br>**S405**<br>**F** |  | A59T, E179K,<br>A620T, R683K,<br>P1305L,<br>L1337F,<br>G1473A, I1474V | N396S, **E461K**,<br>D776N, N896S,<br>L941P |  | V27I,<br>L171P,<br>L248V,<br>V341E,<br>L592F,<br>E1020Q |
| 4 | 1 | E116<br>D,<br>**E266**<br>**D,**<br>**V488I** |  | A59T, E179K,<br>E394K, S484A,<br>R683K, L1337F,<br>G1473A, I1474V | V104F, N396S,<br>**R673Q**, D776N,<br>N896S, L941P | I285T<br>,<br>S405<br>T | P19L,<br>G75R,<br>N937K,<br>E1020Q,<br>F1032L,<br>S1037L |
|  | 2 | **A114**<br>**V,**<br>E116<br>D,<br>**E266**<br>**D,**<br>H283<br>R |  | A59T, T348A,<br>T348I, E394K,<br>S484A, R683K,<br>L1337F, G1473A | V104F, M170V,<br>N396S, D776N,<br>L941P |  | P19L,<br>G75R,<br>N937K,<br>E1020Q,<br>F1032L,<br>S1037L |
| 5 | 1 | E116<br>D,<br>K128<br>T,<br>T172<br>S |  | A59T, E179K,<br>A620T, R683K,<br>V1202F,<br>P1305L,<br>L1337F,<br>G1473A, I1474V | V104F, N199S,<br>H206R, A207V,<br>N396S, N772K,<br>D776N, Q829E,<br>S935*, L941P | I285T<br>,<br>T634I | V27I,<br>L171P,<br>L248V,<br>E1020Q |

